# NaCT (*SLC13A5*) facilitates citrate import and metabolism under nutrient-limited conditions

**DOI:** 10.1101/2021.04.08.439058

**Authors:** Avi Kumar, Thekla Cordes, Anna E. Thalacker-Mercer, Ana M. Pajor, Anne N. Murphy, Christian M. Metallo

## Abstract

Citrate lies at a critical node of metabolism linking tricarboxylic acid metabolism and fatty acid synthesis via acetyl-coenzyme A. Recent studies have linked the sodium citrate transporter (NaCT), encoded by *SLC13A5*, to dysregulated hepatic metabolism and pediatric epilepsy. To examine how NaCT-mediated citrate metabolism contributes to the pathophysiology of these diseases we applied ^13^C isotope tracing to *SLC13A5*-deficient hepatocellular carcinoma (HCC) cell lines and primary rat cortical neurons. Exogenous citrate contributed to intermediary metabolism at appreciable levels only under hypoxic conditions. In the absence of glutamine, citrate supplementation increased *de novo* lipogenesis and growth of HCC cells. Knockout of *SLC13A5* in Huh7 cells compromised citrate uptake and catabolism. Citrate supplementation rescued Huh7 cell viability in response to glutamine deprivation and Zn^2+^ treatment, and these effects were mitigated by NaCT deficiency. Collectively, these findings demonstrate that NaCT-mediated citrate uptake is metabolically important under nutrient limited conditions and may facilitate resistance to metal toxicity.

## Introduction

Citrate lies at a critical node of intermediary metabolism, serving as a substrate for biosynthesis, acetylation, and the regeneration of NAD(P)H. Within mitochondria, citrate is synthesized by citrate synthase and metabolized in the tricarboxylic acid (TCA) cycle to support bioenergetics. Citrate is also exported to the cytosol via the mitochondrial citrate transporter (SLC25A1/CTP) and metabolized by ATP-citrate lyase (ACLY) to generate acetyl-coenzyme A (AcCoA) and downstream metabolic processes, including lipid biosynthesis and acetylation (Wakil and Abu-Elheiga, 2009). Although mitochondrial production of citrate is the major source for most cells, plasma concentrations are relatively high (Costello and Franklin, 2016). In addition, dysregulation of plasma citrate homeostasis has pathophysiological consequences including impaired blood clotting and bone disorders (Costello and Franklin, 2016; Zuckerman and Assimos, 2009). The uptake and utilization of exogenous citrate by cells has therefore garnered increasing interest (Bhutia et al., 2017; Costello and Franklin, 1991, 2016).

A plasma membrane specific variant of *SLC25A1*, pmCIC, allows for import and catabolism of extracellular citrate in prostate cancer cells (Mycielska et al., 2018). Further, several members of the SLC13 sodium sulfate/carboxylate symporter gene family are capable of importing citrate into cells in a sodium-coupled mechanism (Pajor, 2006, 2014). Both *SLC13A2*, primarily expressed in the kidney and small intestine, and *SLC13A3*, widely expressed across tissues, are capable of importing citrate, although with significantly lower affinity compared to dicarboxylic TCA intermediates (Pajor, 2006, 2014). Liver and brain tissue express the sodium-dependent citrate transporter (NaCT; also known as mINDY) encoded by *SLC13A5*, which preferentially transports citrate across the plasma membrane (Inoue et al., 2002; Pajor, 1996).

*SLC13A5* function has been linked to hepatic glucose and fatty acid metabolism and development of NaCT inhibitors has garnered therapeutic interest (Huard et al., 2015; Sauer et al., 2021). Deletion of *Slc13a5* is protective against high-fat diet-induced insulin resistance and attenuates hepatic gluconeogenesis and lipogenesis (Birkenfeld et al., 2011). Further, pharmacological inhibition of NaCT reduced hepatic lipid accumulation in mice fed a high-fat diet (Huard et al., 2015). Additionally, hepatic *SLC13A5* expression correlated positively with both body and liver fat in a cohort of non-alcoholic fatty liver disease (NAFLD) patients (von Loeffelholz et al., 2017). Functional impacts of NaCT inhibition or knockdown have been proposed in cells, though findings are not necessarily dependent on the presence of extracellular citrate (Li et al., 2017; Phokrai et al., 2018; Poolsri et al., 2018). Citrate transport may also influence AcCoA metabolism and ionic homeostasis in the nervous system, as loss-of-function mutations in *SLC13A5* have been linked to pediatric epilepsy, Kohlschütter-Tönz syndrome, and other brain disorders (Hardies et al., 2015; Klotz et al., 2016; Matricardi et al., 2020; Schossig et al., 2017; Thevenon et al., 2014). Notably, citrate levels were significantly increased in plasma and cerebrospinal fluid (CSF) in such epilepsy patients (Bainbridge et al., 2017). Mice deficient in this transporter accumulated citrate in CSF while plasma abundances were not affected; moreover, *Slc13a5*^-/-^ mice exhibited an increased propensity for seizures as well as impaired neuronal function (Henke et al., 2020). While citrate’s function in chelating metal cations has been hypothesized to drive this pathophysiology (Bhutia et al., 2017; Glusker, 1980), the mechanism(s) through which *SLC13A5*-deficiency drives pathogenesis in mammalian cells warrants further study.

Here, we have applied mass spectrometry and metabolic flux approaches to genetically engineered HCC cell lines and primary cortical neurons to decipher the impact of extracellular citrate import on metabolism and cell viability. *SLC13A5* was expressed in both cell types, and exogenous citrate was imported and metabolized to fatty acids and TCA cycle intermediates. However, citrate only contributed appreciably to these pathways under hypoxic conditions where pyruvate dehydrogenase (PDH) flux is reduced. Under these conditions citrate was predominantly catabolized in the cytosol to support AcCoA generation. We also used CRISPR/Cas9 to generate *SLC13A5*-deficient HCC cell lines, which lacked the ability to transport and metabolize exogenous citrate. In addition, we observed that *SLC13A5* expression was required to increase the growth of hypoxic HCC cells under glutamine-deprived conditions. Finally, NaCT function was also necessary to allow for citrate-mediated protection against Zn^2+^ toxicity. Collectively, our study highlights the biological roles of NaCT and citrate metabolism in mammalian cells, emphasizing the importance of metabolic stress in observing significant phenotypes.

## Results

### Extracellular citrate uptake is tissue specific

Recent findings have highlighted the importance of circulating TCA intermediates as metabolic substrates or regulators of tissue function (Mills et al., 2018). We quantified levels of TCA intermediates in human plasma and observed that the concentration of citrate was 108 ± 23 µM while other TCA cycle intermediates were 10-fold less abundant (all < 10 µM) (Fig. 1A). Although citrate is not present in typical culture media such as DMEM, RPMI, or OptiMEM, we found that complete medium containing 10% FBS contained 16 ± 5 µM citrate, significantly lower than that observed in human plasma (Fig. 1B).

**Figure 1.**
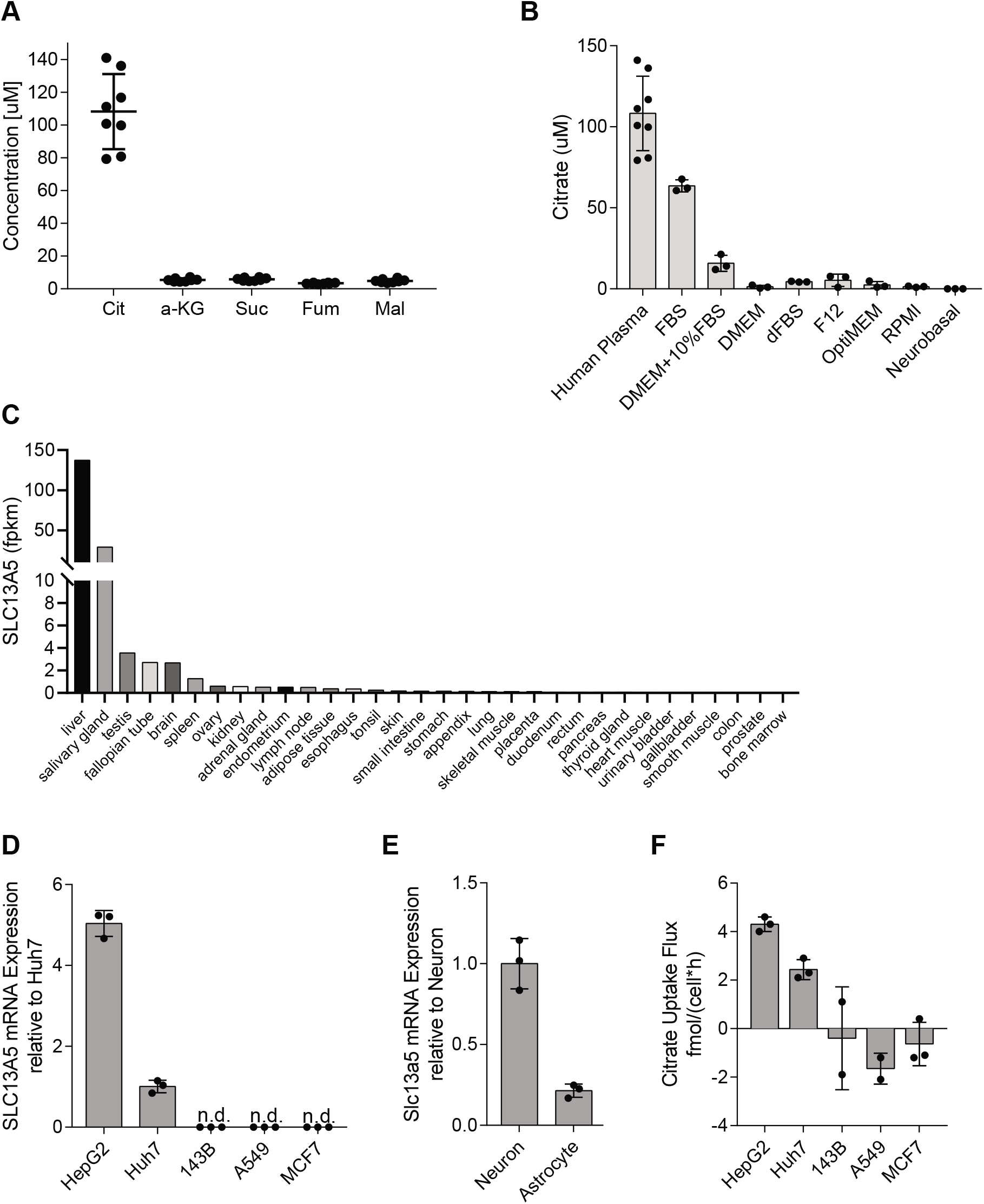
Extracellular citrate uptake is tissue specific. (A) Plasma concentration of TCA intermediates in humans (n=8). (B) Citrate concentration in cell culture medias (n=3), human plasma (n=8) same as in panel 1A. (C) *SLC13A5* mRNA expression in human tissues, from GTEx (Ardlie et al., 2015). (D) *SLC13A5* mRNA expression in cancer cells grown in normoxia relative to Huh7s (n=3). (E) *Slc13a5* mRNA expression in primary rat neuron and astrocyte cells in normoxia relative to neurons (n=3). (F) Net citrate uptake flux in cancer cell lines over 48 hours (n=3). All data are presented as means ± SD. All data are representative of three sample replicates.

To identify cell types that might readily consume extracellular citrate, we first utilized a publicly available transcriptomics dataset from the Genotype-Tissue Expression (GTEx) project and compared *SLC13A5* expression across 32 human tissues (Ardlie et al., 2015). *SLC13A5* transcripts were highest in liver, brain, reproductive tissues, and salivary gland (Fig. 1C). Next, we quantified the expression of *SLC13A5* in a panel of different cells from different tissues of origin. Consistent with published data (Ardlie et al., 2015), we found that the HCC cell lines HepG2 and Huh7 as well as primary cortical neurons express detectable *SLC13A5* mRNA while little was detected in other cell lines (Figs. 1D, E). We next cultured cells in DMEM supplemented with 500 µM extracellular citrate for 48 hours and quantified uptake flux. Only the HCC cell lines exhibited net uptake of citrate, which correlated with *SLC13A5* transcription (Fig. 1F).

### Citrate dilutes central carbon metabolism in HCC cells and neurons in hypoxia

The above results indicate that HCC cells import exogenous citrate. To quantify the contribution of citrate to central carbon metabolism in more detail, we cultured HepG2 and Huh7 cells in medium containing uniformly ^13^C labeled ([U-^13^C_5_]) glutamine and quantified metabolite abundance and isotope enrichment (Fig. 2A). Studies were performed under normoxic (21% oxygen) or hypoxic (1% oxygen) conditions for 48 hours, as hypoxia is known to potently reduce citrate abundances due to decreased PDH flux (Metallo et al., 2012; Wise et al., 2011). Indeed, TCA intermediate abundances were significantly decreased in hypoxia, with citrate being the most decreased of those measured (Fig. S1A). As expected, labeling from [U-^13^C_5_]glutamine indicated that cells increased the contribution of reductive carboxylation to synthesis of aspartate and fatty acids, as well as glutamine anaplerosis under hypoxia (Figs. S1B-D). To indirectly determine whether exogenous citrate was metabolized in cells we cultured Huh7 and HepG2 cells with and without 500 µM unlabeled (^12^C) citrate in medium containing [U-^13^C_5_]glutamine or [U-^13^C_6_]glucose, respectively. Citrate addition resulted in a dilution of glutamine’s contribution to TCA cycle anaplerosis (Figs. 2B, S1E) and *de novo* lipogenesis through reductive carboxylation in HCC cell lines (Figs. 2C, D). We performed similar studies using primary rat cortical neurons, although [U-^13^C_6_]glucose was used given the higher enrichment obtained in differentiated neurons (Divakaruni et al., 2017). Although labeling of the intracellular citrate pool was significantly diluted, glucose-derived TCA labeling was not impacted (Fig. 2E). On the other hand, we observed a significant impact on the contribution of glucose to lipogenic AcCoA using ISA modeling (Fig. 2F). Our findings suggest that extracellular citrate metabolically contributes as an anaplerotic and/or lipogenic substrate in low oxygen conditions, and further highlight the need for some metabolic stress to observe a significant contribution.

**Figure 2.**
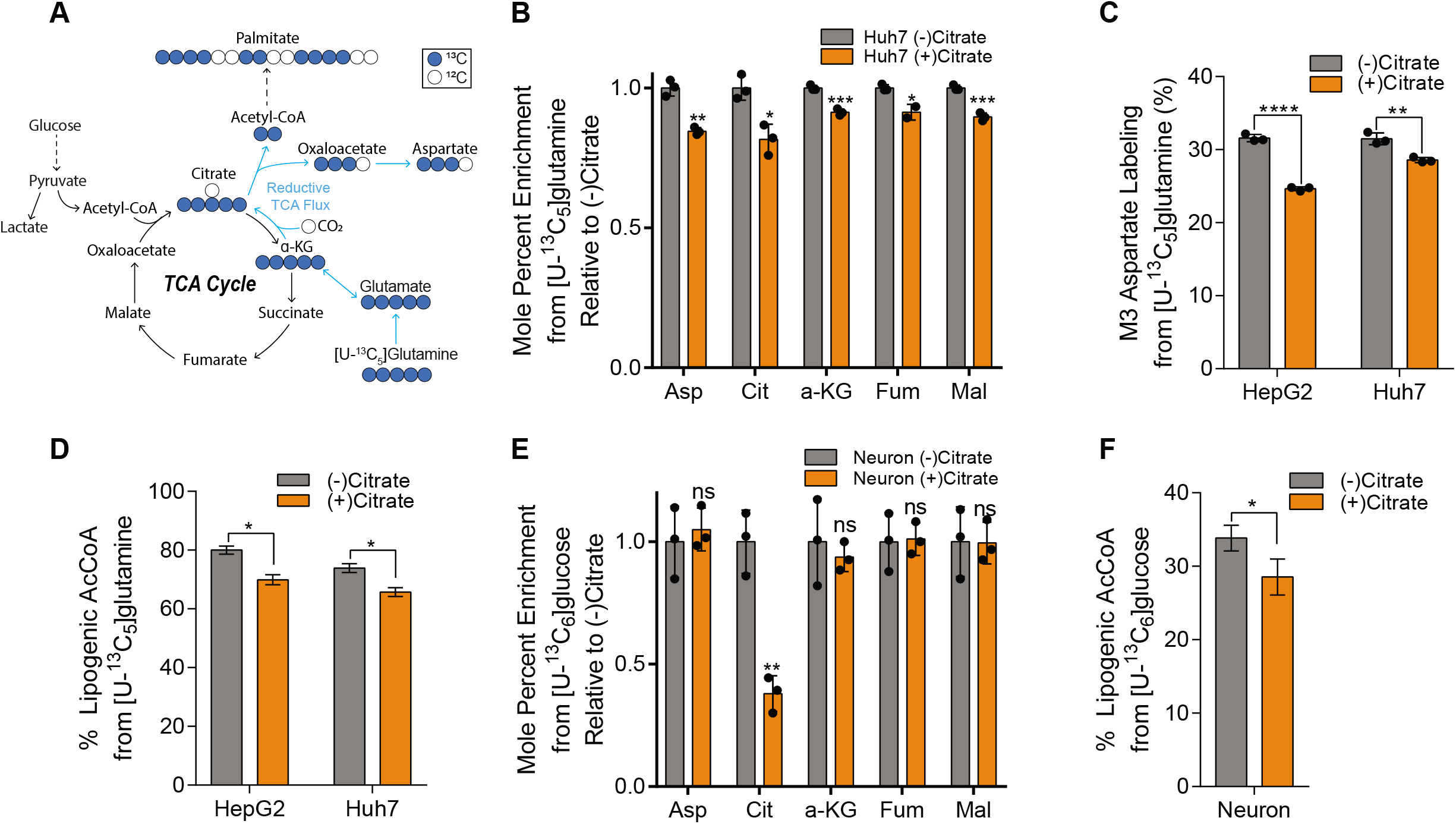
Citrate dilutes central carbon pathway labeling in hepatocellular carcinoma and neuronal cells in hypoxia. (A) Atom transition map depicting catabolism of [U-^13^C_5_]glutamine. Closed circles indicate ^13^C carbon, open circles indicate ^12^C carbon. (B) Mole percent enrichment of TCA intermediates from [U-^13^C_5_]glutamine in Huh7 cells grown in hypoxia +/- 500 µM citrate for 48 hours, relative to (-) citrate (n=3). (C) Percent labeling of M3 aspartate from [U-^13^C_5_]glutamine in HepG2 and Huh7 cells grown in hypoxia +/- 500 µM citrate for 48 hours (n=3). (D) Percent of lipogenic acetyl-CoA contributed by [U-^13^C_5_]glutamine in HepG2 and Huh7 cells grown in hypoxia +/- 500 µM citrate for 48 hours (n=3). (E) Mole percent enrichment of TCA intermediates from [U-^13^C_6_]glucose in primary rat neuron cells grown in 3% oxygen +/- 500 µM citrate for 48 hours, relative to (-) citrate (n=3). (F) Percent of lipogenic acetyl-CoA contributed by [U-^13^C_6_]glucose in primary rat neuron cells grown in 3% oxygen +/- 500 µM citrate for 48 hours (n=3). In (B,C,E) data are plotted as mean ± SD. Statistical significance is relative to (-) citrate as determined by two-sided Student’s t-test with *, P value < 0.05; **, P value < 0.01; ***, P value < 0.001, ****, P value < 0.0001. In (D,F) data are plotted as mean ± 95% confidence interval (CI). Statistical significance by non-overlapping confidence intervals, *. Unless indicated, all data represent biological triplicates. Data shown are from one of at least two separate experiments. See also Figure S1.

To more directly quantify how extracellular citrate is metabolized in *SLC13A5*-expressing cells, we cultured the above cell types in growth medium supplemented with 500 µM [2,4-^13^C_2_]citrate. Cells were rinsed 2X in NaCl prior to extraction to ensure accurate analysis of intracellular pools. In HepG2 and Huh7 cells, extracellular citrate contributed significantly to TCA labeling, with ∼14% labeling on citrate and ∼5% enrichment on downstream intermediates in normoxic cells (Fig. 3A). Under hypoxia, enrichment of intracellular citrate and other TCA intermediates was significantly elevated (Fig. 3A). Intracellular citrate was also highly enriched by [2,4-^13^C_2_]citrate in neonatal rat cortical neurons, though detectable labeling was only observed on alpha-ketoglutarate (Fig. 3B). Furthermore, no changes in enrichment were observed when comparing normoxic to 3% oxygen. These differences are likely due to the reduced anaplerotic/biosynthetic needs of post-mitotic neurons. However, our results confirm that extracellular citrate is transported into these *SLC13A5-*expressing cell types. Importantly, we were unable to detect significant isotope enrichment on TCA intermediates in other cell types tested that lack *SLC13A5* expression, suggesting this transporter is important for citrate import and metabolism (Fig. S2A).

**Figure 3.**
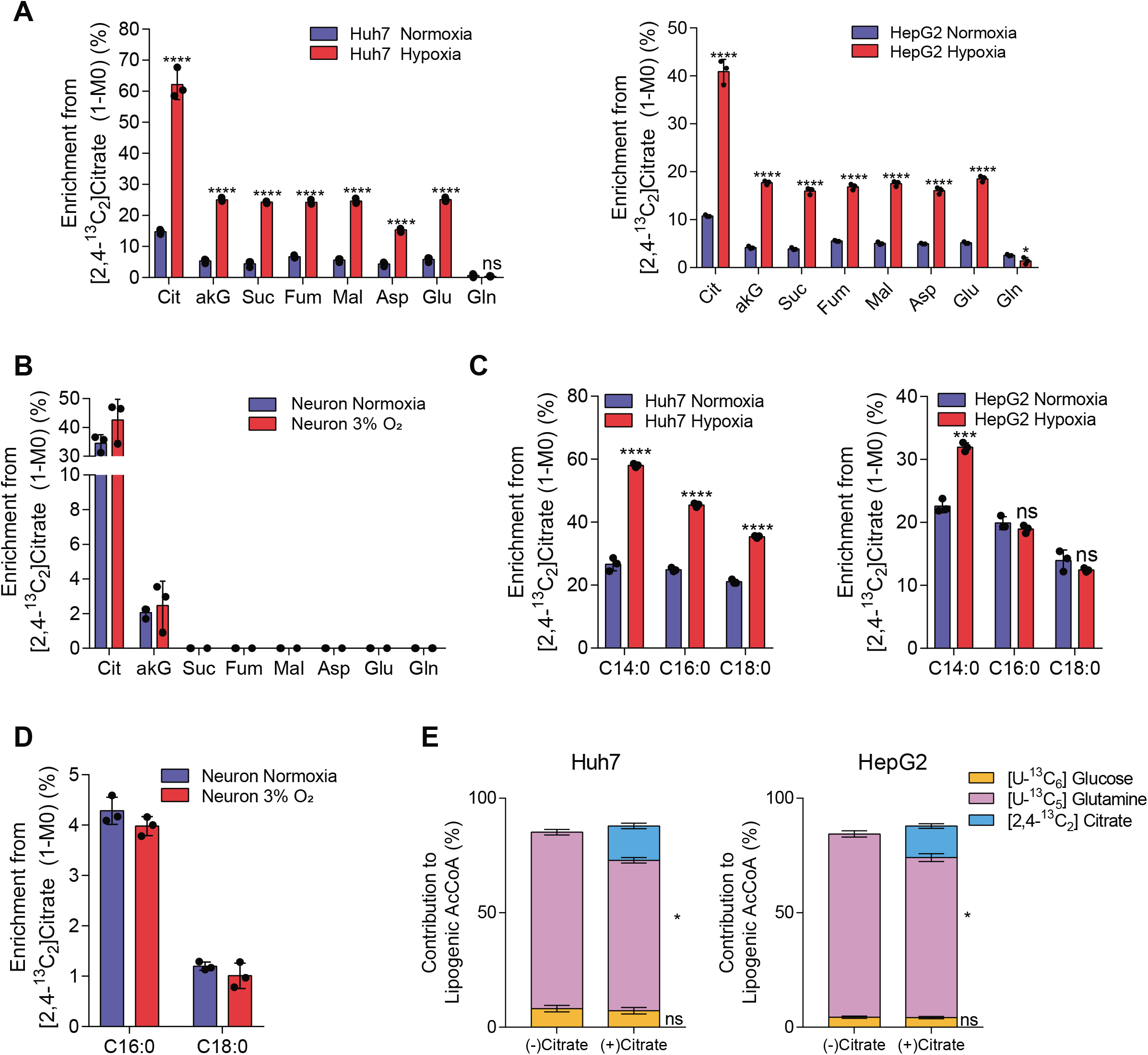
Exogenous citrate is metabolized for TCA anaplerosis and fatty acid synthesis. (A) Enrichment (1-M0) of TCA intermediates from [2,4-^13^C_2_]citrate in Huh7 (left) and HepG2 (right) cells grown in normoxia or hypoxia for 48 hours (n=3). (B) Enrichment (1-M0) of TCA intermediates from [2,4-^13^C_2_]citrate in primary rat neuron cells grown in normoxia or 3% oxygen for 48 hours (n=3). (C) Enrichment (1-M0) of fatty acids from [2,4-^13^C_2_]citrate in Huh7 (left) and HepG2 (right) cells grown in normoxia or hypoxia for 48 hours (n=3). (D) Enrichment (1-M0) of fatty acids from [2,4-^13^C_2_]citrate in primary rat neuron cells grown in normoxia or 3% oxygen for 48 hours (n=3). (E) Contribution of [U-^13^C_6_]glucose, [U-^13^C_5_]glutamine and [2,4-^13^C_2_]citrate in the presence or absence of 500 µM unlabeled citrate in hypoxia in Huh7 (left) and HepG2 (right) cells (n=3). In (A-D) data are plotted as mean ± SD. Statistical significance is relative to normoxia as determined by two-sided Student’s t-test with *, P value < 0.05; **, P value < 0.01; ***, P value < 0.001, ****, P value < 0.0001. Unless indicated, all data represent biological triplicates. In (E) data are plotted as mean ± 95% confidence interval (CI). Statistical significance by non-overlapping confidence intervals, *. Data shown are from one of at least two separate experiments. See also Figure S2.

Next, we examined whether extracellular citrate was metabolized to AcCoA by quantifying enrichment of fatty acids on cells cultured with the isotope tracers noted above. We detected significant enrichment of palmitate and stearate from [2,4-^13^C_2_]citrate, demonstrating that HCC cell lines and neurons generate AcCoA from exogenous citrate (Figs. 3C, D). No enrichment of palmitate was observed in other cell types that lack *SLC13A5* expression (Fig. S2B). We then compared the relative contributions of [U-^13^C_5_]glucose, [U-^13^C_5_]glutamine, and [2,4-^13^C_2_]citrate to the lipogenic AcCoA fueling palmitate synthesis under hypoxia (Fig. 3E). As expected, glutamine is the primary source of lipogenic carbon under these conditions. However, when citrate was present in the medium, HCC cells actively metabolized this substrate and the contribution of exogenous citrate to palmitate was greater than that of glucose, though glutamine remained the major lipogenic substrate in these hypoxic conditions. Notably, extracellular citrate did not alter the rate of *de novo* lipogenesis (Fig. S2C). Thus, exogenous citrate is taken up by *SLC13A5* expressing cell types and is metabolized to fuel TCA cycle metabolism and lipogenesis, thereby diluting the corresponding glutamine contribution in low oxygen conditions. These findings are reminiscent of the usage of acetate under hypoxia in cells expressing ACSS2 (Kamphorst et al., 2014).

### Extracellular citrate is primarily catabolized in the cytosol

The [2,4-^13^C_2_]citrate tracer enables more detailed insights into the compartment specific catabolism of extracellular citrate when considering isotopologue distributions in more detail. In hypoxic, proliferating cancer cells, TCA intermediates are predominantly obtained from glutaminolysis (M4 labeling from [U-^13^C_5_]glutamine) or reductive carboxylation (M3 labeling from [U-^13^C_5_]glutamine), with TCA (re)cycling contributing to a lesser degree (Fan et al., 2013; Metallo et al., 2012). Therefore, TCA substrates in such cells are mostly present for one pass of the cycle. When [2,4-^13^C_2_]citrate is catabolized directly in the cytosol by ATP-citrate lyase (ACLY), M1 acetyl-CoA and M1 oxaloacetate are formed which further yield M1 TCA intermediates via the malate-aspartate shuttle (Fig. 4A). Alternatively, if citrate is imported into the mitochondria directly by SLC25A1, as has been shown to occur in anchorage-independent cultures (Jiang et al., 2016a, 2016b), [2,4-^13^C_2_]citrate will generate M2 isotopologues of TCA intermediates (Fig. 4B). When culturing HCC cell lines in the presence of 500 µM [2,4-^13^C_2_]citrate for 48 hours in either normoxia or hypoxia, we found that the M1 isotopologue of aspartate, malate and fumarate was more abundant than the M2 isotopologue, and the ratio of M1/M2 labeling increased for these metabolites in hypoxic conditions (Fig. 4C). On the other hand, M2 alpha-ketoglutarate relative abundance was similar to or greater than M1 labeling, which indicates that a high exchange flux exists for the cytosolic aconitase (ACO1) and isocitrate dehydrogenase (IDH1) reactions (Figs. 4A, D). Collectively, these results suggest that exogenous citrate is primarily metabolized in the cytosol by ACLY and ACO1/IDH1 under the conditions tested (Figs. 4C, D).

**Figure 4.**
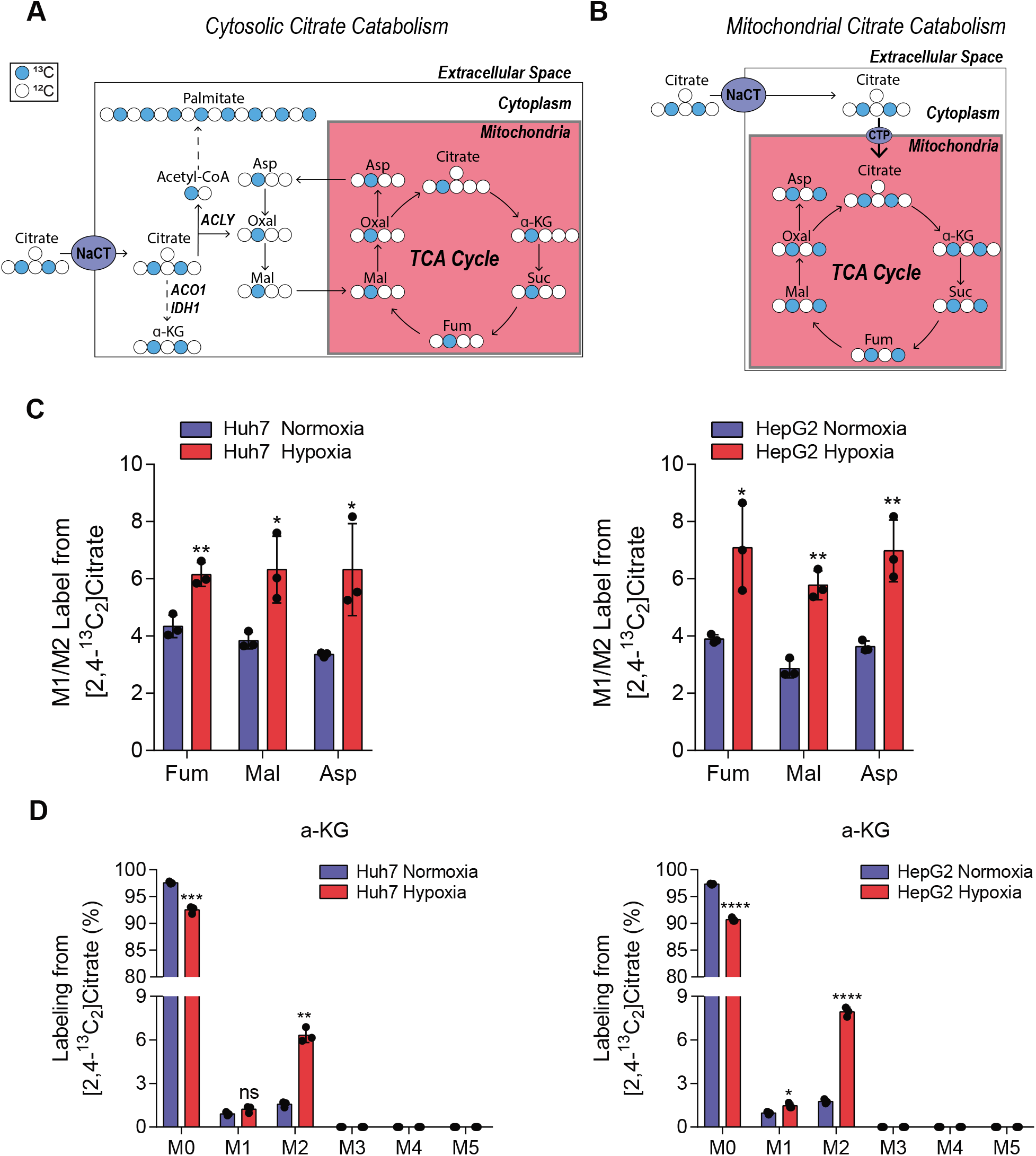
Extracellular citrate is primarily catabolized in the cytosol. (A) Schematic of extracellular [2,4-^13^C_2_]citrate catabolism in the cytosol. Closed circles indicate ^13^C carbon, open circles indicate ^12^C carbon. (B) Schematic of extracellular [2,4-^13^C_2_]citrate catabolism in the mitochondria. Closed circles indicate ^13^C carbon, open circles indicate ^12^C carbon. (C) Ratio of the relative abundance of M1/M2 TCA intermediates from 500 µM [2,4-^13^C_2_]citrate in Huh7 (left) and HepG2 (right) cells over 48 hours in hypoxia (n=3). (D) Mass isotopomer distribution of aKG from 500 µM [2,4-^13^C_2_]citrate in Huh7 (left) and HepG2 (right) cells over 48 hours in hypoxia (n=3). In all graphs data are plotted as mean ± SD. Statistical significance is relative to normoxia as determined by two-sided Student’s t-test with *, P value < 0.05; **, P value < 0.01; ***, P value < 0.001, ****, P value < 0.0001. Unless indicated, all data represent biological triplicates. Data shown are from one of at least two separate experiments.

### NaCT coordinates citrate import and metabolism in hepatocellular carcinoma cells

To directly interrogate the function of NaCT with respect to citrate metabolism, we engineered *SLC13A5*-deficient knockout (KO) HepG2 and Huh7 cells using CRISPR/Cas9 and compared their metabolic phenotype to cells expressing a non-targeting control (NTC) sgRNA. Two clones were tested for each KO cell line. *SLC13A5* KO clones were validated by sequencing the region of interest (Fig. S3). In both cell lines, knockout of *SLC13A5* resulted in reduced citrate uptake flux when the cells were cultured in growth medium supplemented with 500 µM exogenous citrate (Figs. 5A, S4A), suggesting these cells lacked NaCT activity. To further verify the functional impact of NaCT deficiency we quantified citrate transport using [1,5-^14^C]citrate (Pajor et al., 2016). Sodium induced citrate uptake was significantly reduced in the *SLC13A5* KO cells (Fig. 5B). Notably, NaCT deficiency induced no significant metabolic phenotype in cells grown in the absence of extracellular citrate, as fatty acid synthesis rates and metabolite abundances were not altered in the knockout clones (Figs. S4B, C). Additionally, although previous studies have found that *SLC13A5* knockdown alone induced a reduction in the expression of fatty acid synthesis gene expression in hepatocellular carcinoma cells (Li et al., 2017), we observed no impact on ACLY expression in HepG2 or Huh7 *SLC13A5* KO cells (Fig. S4D). These data indicate that the citrate transporter induces a metabolic change that is selective to citrate uptake.

**Figure 5.**
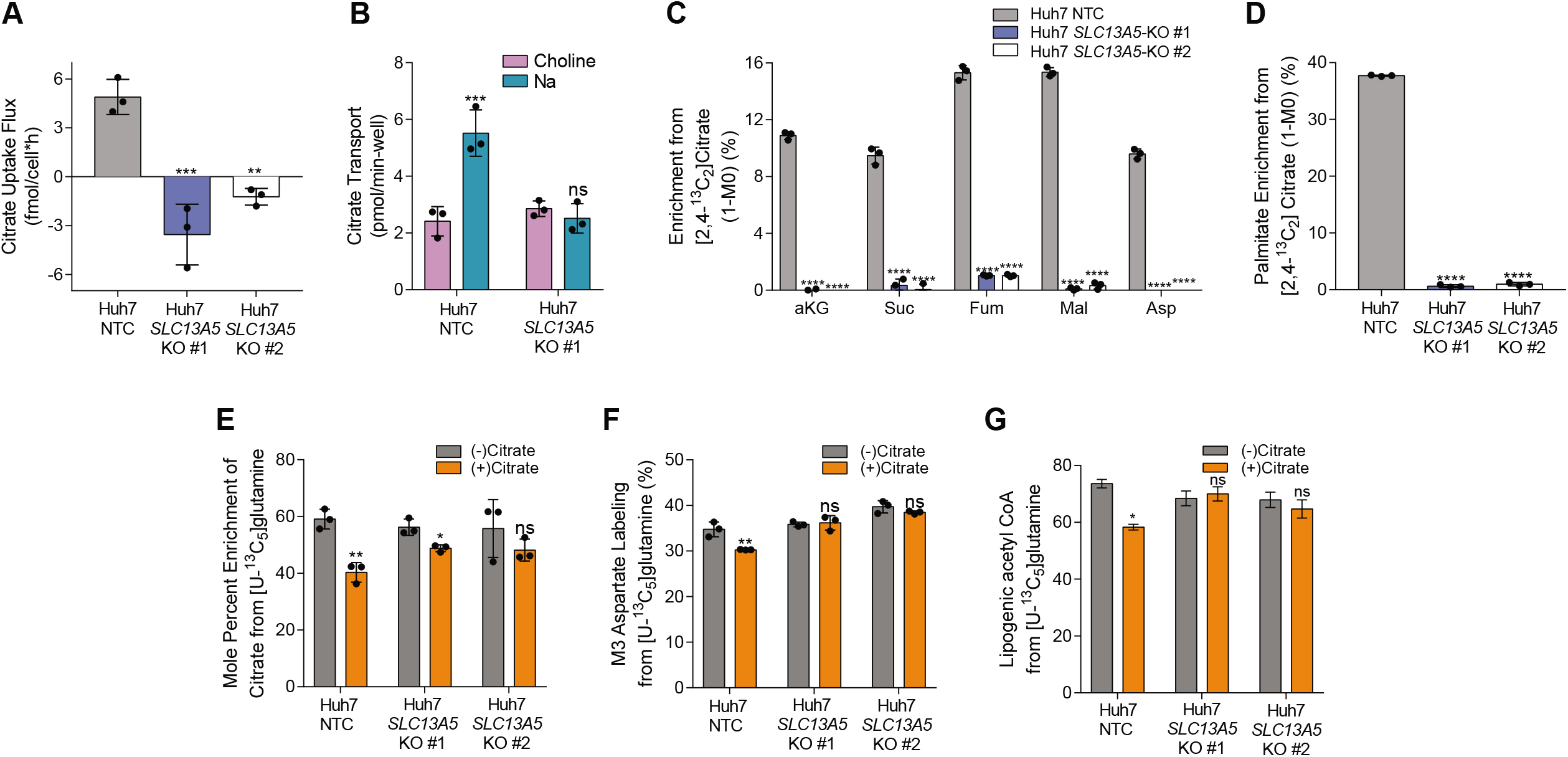
NaCT coordinates citrate import and metabolism in hepatocellular carcinoma cells. (A) Citrate uptake flux over 48 hours in hypoxia in Huh7 NTC and *SLC13A5*-KO cells (n=3). (B) Citrate transport in Huh7 NTC and *SLC13A5*-KO #1 cells (n=3). (C) Enrichment (1-M0) of TCA intermediates from [2,4-^13^C_2_]citrate in Huh7 NTC and *SLC13A5*-KO cells grown in hypoxia for 48 hours (n=3). (D) Enrichment (1-M0) of palmitate from [2,4-^13^C_2_]citrate in Huh7 NTC and *SLC13A5*-KO cells grown in hypoxia for 48 hours (n=3). (E) Mole percent enrichment of citrate from [U-^13^C_5_]glutamine in Huh7 NTC and *SLC13A5*-KO cells +/- 500 µM citrate grown in hypoxia for 48 hours (n=3). (F) Percent labeling of M3 aspartate from [U-^13^C_5_]glutamine in Huh7 NTC and *SLC13A5*-KO cells +/- 500 µM citrate grown in hypoxia for 48 hours (n=3). (G) Percent of lipogenic acetyl-CoA contributed by [U-^13^C_5_]glutamine in Huh7 NTC and *SLC13A5*-KO cells +/- 500 µM citrate grown in hypoxia for 48 hours (n=3). In all graphs data are plotted as mean ± SD. Statistical significance is relative to NTC as determined by One-way ANOVA w/ Dunnet’s method for multiple comparisons (A,C,D) or relative to (-) citrate as determined by two-sided Student’s t-test (B,E,F) with *, P value < 0.05; **, P value < 0.01; ***, P value < 0.001, ****, P value < 0.0001. In (G) data are plotted as mean ± 95% confidence interval (CI). Statistical significance by non-overlapping confidence intervals, *. Unless indicated, all data represent biological triplicates. See also Figure S3 and S4. (C-G) Data shown are from one of at least two separate experiments.

To elucidate how impaired NaCT activity influences citrate catabolism, we cultured each cell line for 48 hours in the presence of 500 µM [2,4-^13^C_2_]citrate hypoxic conditions. Huh7 *SLC13A5* KO cells had no measurable enrichment from [2,4-^13^C_2_]citrate on TCA cycle intermediates or fatty acids (Figs. 5C, D). A reduction in enrichment was also observed with HepG2 *SLC13A5* KO cells (Fig. S4E, F). Our data confirms that NaCT is the primary importer of extracellular citrate in the tested HCC cell lines. Next, we cultured NaCT*-*deficient cells in the presence or absence of exogenous ^12^C citrate in [U-^13^C_5_]glutamine tracer medium and quantified enrichment on TCA cycle intermediates and fatty acids compared to control cells. When unlabeled citrate was present in the media, we only observed dilution of [U-^13^C_5_]glutamine on TCA cycle intermediates in *SLC13A5-*expressing cells (Figs. 5E, S4G). Furthermore, we observed no dilution in isotopologues downstream of reductive carboxylation (i.e. M3 aspartate) or glutamine-derived lipogenic acetyl-CoA with extracellular unlabeled citrate addition in the Huh7 *SLC13A5* KO cells (Figs. 5F, G). Thus, our results indicate that NaCT is required for the import and metabolism of extracellular citrate, in HCC cells.

### NaCT facilitates growth under nutrient stress and resistance to zinc toxicity

Next we hypothesized that citrate import and catabolism could fulfill the metabolic role of glutamine, which serves as a key substrate for anaplerosis and acetyl-CoA synthesis under hypoxia. We cultured the above HCC cell lines in glutamine-deficient media containing [U-^13^C_5_]glucose supplemented with exogenous citrate. Notably, addition of citrate in the absence of glutamine significantly increased fatty acid synthesis in control cells (Fig. 6A). Furthermore, citrate supplementation generally decreased enrichment of TCA intermediates and associated amino acids from [U-^13^C_5_]glucose in NTC cells (Fig. 6B), no change was observed in NaCT-deficient cell lines. Next, we cultured *SLC13A5* expressing Huh7 cells with [2,4-^13^C_2_]citrate and found that in the absence of glutamine, labeling from ^13^C citrate on metabolites associated with glutamine synthesis increased significantly (Fig. 6C). Further, ^13^C citrate did not label glutamine in cells lacking NaCT or in glutamine replete medium (Fig. S5A). These data indicate that NaCT facilitates citrate-mediated anaplerosis under nutrient limited conditions.

**Figure 6.**
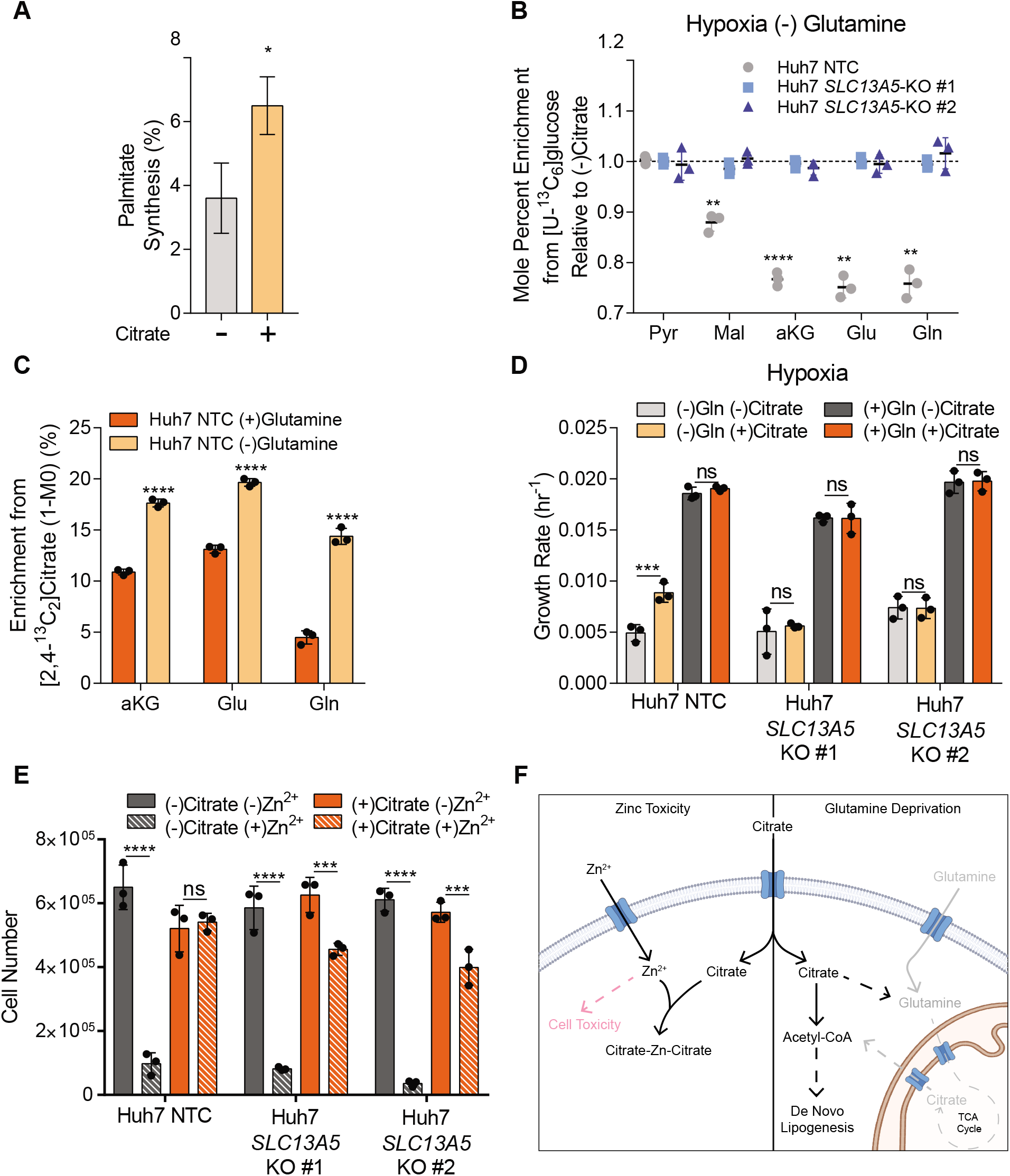
NaCT facilitates growth under nutrient stress and resistance to zinc toxicity. (A) Palmitate synthesis in Huh7 NTC cells grown in high glucose DMEM without glutamine +/- 500 µM citrate in hypoxia for 48 hours (n=3). (B) Mole percent enrichment of metabolites from [U-^13^C_6_]glucose in Huh7 *SLC13A5*-KO cells grown without glutamine in hypoxia +/- 500 µM citrate for 48 hours, relative to (-) citrate (n=3). (C) Enrichment (1-M0) of TCA intermediates from [2,4-^13^C_2_]citrate in Huh7 NTC cells grown in hypoxia +/- 4 mM glutamine for 48 hours (n=3). (D) Growth rates of Huh7 NTC and *SLC13A5*-KO cells grown in high glucose DMEM +/- 4mM glutamine +/- 500 µM citrate in hypoxia for 4 days. (E) Cell numbers of Huh7 NTC and *SLC13A5*-KO cells grown in high glucose DMEM +/- 5mM citrate +/- 300 µM ZnCl_2_ 24 hours after zinc treatment in hypoxia. (F) Schematic depicting extracellular citrate utilization during cellular stress. Gray indicates removed metabolites, pink indicates cytotoxic. In (A) data are plotted as mean ± 95% confidence interval (CI). Statistical significance by non-overlapping confidence intervals, *. In (B-E) data are plotted as mean ± SD. Statistical significance is determined by two-sided Student’s t-test relative to (-) citrate (B), (+) glutamine (C); or determined by Two-way ANOVA w/ Tukey’s method for multiple comparisons relative to (-) citrate (D) or (-) Zn^2+^ conditions (E) with *, P value < 0.05; **, P value < 0.01; ***, P value < 0.001, ****, P value < 0.0001. Unless indicated, all data represent biological triplicates. See also Figure S5. Data shown are from one of at least two separate experiments.

Since glutamine is critical for numerous biological processes, including anaplerosis and lipogenesis, we next tested whether NaCT-mediated citrate uptake impacted cell proliferation upon glutamine deprivation. We observed that exogenous citrate addition increased the growth rate of the Huh7 NTC cells by 80% in glutamine depleted media under hypoxic conditions (Fig. 6D). Notably, no growth alteration was observed with citrate addition in glutamine rich media (Fig. 6D) or normoxia (Fig. S5B). Importantly, citrate supplementation did not increase proliferation of NaCT-deficient cell lines (Figs. 6D, S5B). This data indicates that NaCT-mediated citrate transport supports metabolism under nutrient limited conditions. We next tested whether NaCT-mediated citrate transport influences resistance to metal toxicity against ions such as zinc (Zn^2+^). Regions of the brain including the hippocampus and cerebral cortex are Zn^2+^ enriched (Assaf and Chung, 1984). Under pathological conditions including epilepsy, ischemia, and traumatic brain injury, cerebral Zn^2+^ levels increase and may cause neuronal cell death (Assaf and Chung, 1984; Frederickson et al., 1983; Weiss et al., 2000). Intracellular citrate has been suggested to be a protective chelator against Zn^2+^ in rat cortical neurons (Sul et al., 2016), so we employed the above HCC cell model to determine if NaCT plays a role in this process. We hypothesized that NaCT-mediated citrate uptake facilitates protection against Zn^2+^ toxicity. We cultured hypoxic Huh7 NTC and NaCT-deficient cells in the presence or absence of citrate and ZnCl_2_ for 24 hours and quantified cell viability. While citrate supplementation protected Huh7 NTC against Zn^2+^ toxicity, cells lacking NaCT showed reduced viability (Fig 6E). Thus, our data highlights that extracellular citrate import via NaCT facilitates protection against nutrient deprivation as well as Zn^2+^ toxicity in human cells (Fig. 6F).

## Discussion

Cells take up diverse nutrients from the extracellular microenvironment, and each metabolite may serve a distinct purpose or function. Nutrient transport is selective and regulated, in part, through cell-specific expression of transporters such as NaCT, which is restricted to liver and neural tissue. Here we investigated how NaCT influences citrate metabolism in HCC cell lines and primary rat cortical neurons. We observed that NaCT-mediated citrate import contributes to cytosolic acetyl-CoA pools in HCC cells and neurons under distinct conditions. Hypoxic HCC cells metabolized extracellular citrate to fatty acids and TCA intermediates in a NaCT-dependent manner. When glutamine (and other nutrients) became limited, citrate supported the growth and lipid synthesis of HCC cell lines expressing NaCT. Furthermore, citrate supplementation enhanced the viability of HCC cell lines exposed to Zn^2+^ toxicity, but this protective effect was mitigated in *SLC13A5* KO cells that lacked NaCT expression.

Many cancer cell types increase glutamine utilization to support the TCA cycle and fatty acid biosynthesis *in vitro* (Metallo et al., 2012; Mullen et al., 2012); however reproducing this phenotype *in vivo* has generated mixed results (Davidson et al., 2016; Le et al., 2012; Son et al., 2013; Tardito et al., 2015). One potential explanation for this discrepancy is the difference in nutrient profile of cell culture media compared to *in vivo* microenvironments (Ackermann and Tardito, 2019; Rossiter et al., 2021; Voorde et al., 2019). Plasma and interstitial fluid contain metabolites that impact cellular metabolism but are routinely excluded from most cell culture medias. For instance, uric acid, a metabolite abundant in human plasma but absent in many cell culture medias, inhibited *de novo* pyrimidine synthesis in cancer cells *in vitro* (Cantor et al., 2017). Previous *in vitro* studies have observed that upregulated reductive glutamine catabolism supports fatty acid biosynthesis and defuses mitochondrial redox stress in hypoxia (Jiang et al., 2016b; Metallo et al., 2012). In our study we found that reductive carboxylation was elevated in hypoxia in HepG2 and Huh7 cells but observed that extracellular citrate addition reduced flux through this pathway. Thus, uptake of extracellular citrate may provide an additional carbon source which is accessible to ACLY and downstream pathways, curtailing the need for other nutrients to support these processes. Similar observations of citrate import and metabolism were made in prostate cancer cells, which express the mitochondrial citrate carrier on their plasma membrane (pmCIC) (Mycielska et al., 2018). On the other hand, we only observed appreciable citrate contributions to biosynthesis under selected conditions (hypoxia and glutamine restriction). These findings collectively suggest that citrate is not a major biosynthetic substrate but serves as a resource to cells when they experience distinct stresses (e.g. ischemia, hypoxia, metal toxicity).

We also demonstrated that *SLC13A5* encoded NaCT is the primary mechanism for citrate import in HCC cells. No major changes in metabolism were observed beyond acetyl-CoA and TCA metabolism, and phenotypic responses were dependent upon expression of functional NaCT as well as citrate supplementation. Importantly, some prior studies observed signaling and growth phenotypes in cells upon knockdown or inhibition of NaCT in the absence of extracellular citrate (Li et al., 2017; Phokrai et al., 2018; Poolsri et al., 2018), but our results indicate that citrate transport is a key function of the *SLC13A5* gene product. Indeed, we also found that citrate uptake by NaCT was protective against Zn^2+^ cytotoxicity in Huh7 cells. This finding mirrors those of previous studies which demonstrated that citrate or pyruvate administration were protective against Zn^2+^ cytotoxicity in neurons (Sul et al., 2016; Yoo et al., 2004). Loss-of-function mutations in *SLC13A5* have been associated with early onset epilepsy, but the role of citrate metabolism in the pathophysiology of these patients is not well understood (Hardies et al., 2015; Klotz et al., 2016; Thevenon et al., 2014). Various mechanisms have been proposed for this phenotype, including the susceptibility of NaCT-deficient neurons to increased synaptic Zn^2+^concentrations induced upon neuronal activation (Sul et al., 2016). Alternatively, others have hypothesized that improper NaCT function leads to increased synaptic citrate concentrations and excessive Zn^2+^chelation which impairs NMDA receptor function and drives neuronal dysfunction (Bhutia et al., 2017). Regardless, our findings should motivate further investigation into the impact of *SLC13A5* mutations on homeostasis of potentially toxic metal cations such as Zn^2+^ and its relation to epilepsy. Collectively, our results demonstrate that NaCT mediates citrate transport and metabolism under conditions of metabolic stress. Our approach also highlights the utility of coordinated studies that involve targeting of disease-related metabolic genes in relevant cell types and deep metabolic profiling using flux analysis.

## Acknowledgments

We thank all members of the Metallo laboratory for support and helpful discussions. This study was supported, in part, by US National Institutes of Health (NIH) grants R01CA234245 (C.M.M.), R21NS104513 (A.N.M., A.M.P.) and a TESS Research Foundation Grant (A.N.M., A.M.P.).

## Author contributions

A.K., T.C. and C.M.M. conceived and designed the study. A.K. performed HCC cell line experiments. A.K. and T.C. generated the knockout cell lines. T.C. and A.N.M performed experiments with cultured primary rat neurons. A.M. P. performed ^14^C citrate uptake experiments. A.T.M. provided human plasma samples. A.K., T.C. and C.M.M. wrote the paper with input from all authors.

## Materials and Methods

### Resource Availability

#### Lead Contact

Further information and requests for resources and reagents should be directed to and will be fulfilled by the Lead Contact, Christian M. Metallo

#### Materials availability

This study did not generate new unique reagents.

#### Data and code availability

The datasets supporting the current study are available from the corresponding author upon request.

### Experimental Model and Subject Details

#### Human Samples

Plasma samples from healthy fasted adults (29-42 years, 5 male, 3 female) were obtained from a clinical cohort recruited from Tompkins County, New York area as described previously (Gheller et al., 2021). Participants were excluded if they had a history of negative or allergic reactions to local anesthetic, used immunosuppressive medications, were prescribed anti-coagulation therapy, were pregnant, had a musculoskeletal disorder, suffered from alcoholism (>11 drinks per week for women and >14 drinks per week for men) or other drug addictions or were acutely ill at the time of participation (Gheller et al., 2019; Riddle et al., 2018). The Cornell University Institutional Review Board approved the protocol and all the subjects provided written informed consent in accordance with the Declaration of Helsinki.

#### Cell Lines

HepG2, Huh7, A549, MCF7, and 143B cells were obtained from ATCC and were incubated at 37C with 5% CO_2_ and cultured using Dulbecco’s Modified Eagle Media (DMEM) with 10% Fetal Bovine Serum and 1% Penicillin-Streptomycin. Cells tested negative for mycoplasma contamination. For hypoxia experiments, cells were maintained in a humidified glove box (Coy) at 5% CO_2_ and either 1% O_2_ (cancer cell lines) or 3% O_2_ (primary cortical neurons). Primary cortical neurons and astrocytes were isolated and cultured in Neurobasal medium (neuron) or DMEM medium supplemented with 10 mM glucose, 0 mM glutamine, and 10% FBS (astrocytes) as described in detail by Cordes et al. 2020 (Cordes et al., 2020). All media were adjusted to pH = 7.3.

### Method Details

#### Cell Proliferation and ^13^C Tracing

Proliferation studies were performed on 12 well plates with an initial cell number of 50,000/well for Huh7s and 100,000/well for HepG2s. Cells were plated in growth media and allowed to adhere for 24 hours before changing to the specified growth media. Cell counts were performed at days 0 and 4 using a hemocytometer.

For Zn^2+^ tolerance studies, cells were plated at 50,000/well and allowed to adhere for 24 hours. They were then placed in the hypoxia chamber for 24 hours before swapping to +/- 5mM citrate media. After changing media, cells were allowed to acclimate for 24 hours before 300 µM ZnCl_2_ in water was spiked into the designated wells. Cells were counted 24 hours after Zn^2+^ treatment using a hemocytometer.

^13^C isotope tracing media was formulated using glucose and glutamine free DMEM 5030 supplemented with either 20 mM [U-^13^C_6_]glucose, 4 mM [U-^13^C_5_]glutamine, or 500 µM [2,4-^13^C_2_]citrate (Cambridge Isotopes) and 10% dialyzed FBS. All studies were performed with a final concentration of 20 mM glucose and 4 mM glutamine. Cultured cells were washed with 1 mL PBS prior to applying tracing media for 24-72 hours as indicated in figure legends. For tracing experiments performed in hypoxia, cancer cells were acclimated to the hypoxia chamber in basal media for 24 hours prior to applying pre-hypoxified tracing media.

#### Isotopomer Spectral Analysis (ISA)

Isotopomer spectral analysis (ISA) was performed to estimate the percent of newly synthesized palmitate as well as the contribution of a tracer of interest to the lipogenic acetyl-CoA pool (Cordes and Metallo, 2019; Young, 2014). Parameters for contribution of ^13^C tracers to lipogenic acetyl-CoA (D value) and percentage of newly synthesized fatty acid (g(t) value) and their 95% confidence intervals are then calculated using best-fit model from INCA MFA software. Experimental fatty acid labeling from [U-^13^C_6_]glucose, [U-^13^C_5_]glutamine, or [2,4-^13^C_2_]citrate after 48 hour trace was compared to simulated labeling using a reaction network where C16:0 is condensation of 8 AcCoA. ISA data plotted as mean ± 95% CI. * indicates statistical significance by non-overlapping confidence intervals.

#### CRISPRcas9 engineered knockout cell lines

*SLC13A5* knockout clones were generated using the strategy described previously (Ran et al., 2013). Briefly, a guide RNA (gRNA) was designed to target the human *SLC13A5* gene using CRISPOR (gRNA sequence: AGGCACAATGAATAGCAGGG) (Concordet and Haeussler, 2018). The gRNA duplex was cloned into lentiCRISPRv2 (Addgene #52961) (Sanjana et al., 2014). HepG2 and Huh7 cells were transfected with the *SLC13A5* specific gRNA to generate pooled knockouts. After puromycin selection, single-cell clones were isolated by diluting the pooled knockout lines at 1 cell/100 µL and plating 100 µL into each well of a 96 well plate. Clones were maintained by exchanging media every 3-5 days. HepG2 clones were cultured in 50% conditioned media to enhance proliferation at early stages of cloning. Conditioned media was generated by collecting the spent media that had been culturing HepG2s after 2 days, spinning down at 300*g* for 5 min, and filtering using a 0.2 micron filter to clear cellular debris. This media was diluted 1:1 with DMEM with 10% FBS 1% penstrep to generate the 50% conditioned media. Clones were validated via Sanger sequencing.

#### Lentivirus Production

One 10cm dish of HEK293FT cells at 60% confluency were transfected with 1.3 µg VSV.G/pMD2.G, 5.3 µg lenti-gag/pol/pCMVR8.2, and 4 µg of the gRNA duplexed lentiCRISPRv2 using 16 µL Lipofectamine 3000 diluted in 0.66 mL of OPTI-MEM. Medium containing viral particles was harvested 48 and 72 hours after transfection, then concentrated by Centricon Plus-20 100,000 NMWL centrifugal ultrafilters, divided into aliquots and frozen at -80°C.

#### Metabolic Flux Analysis

Metabolic fluxes for citrate were calculated by collecting media at time 0 and spent media after 48 hours. Spent media was centrifuged at 300g for 5 min, to remove cell debris. Cell counts were performed at time 0 and after 48 hours as well. To calculate citrate uptake fluxes, citrate was quantified by GC-MS utilizing a standard curve which was run separately.

#### Metabolite Extraction and GC-MS Analysis

At the conclusion of the tracer experiment, media was aspirated. Then, cells were rinsed twice with 0.9% saline solution and lysed with 250 µL ice-cold methanol. After 1 minute, 100 µL water containing 1 µg/ml norvaline was added to each sample and vortexed for one minute. 250 µL chloroform was added to each sample, and all were vortexed again for 1 minute. After centrifugation at 21,130 g for 10 minutes at 4°C, 250 µL of the upper aqueous layer was collected and evaporated under vacuum at 4°C. Then, 250 µL of the upper aqueous layer was collected and evaporated under air at room temperature.

Plasma metabolites were extracted and quantified as follows. For metabolite extraction, 10 µL of each plasma sample was utilized. First, 90 µL of a 9:1 methanol:water mix was added to each sample and the samples were vortexed for 1 minute. After centrifugation at 16,000g for 10 minutes, 90 µL of supernatant was collected, evaporated under vacuum at -4°C and analyzed using GC/MS. Metabolite levels of TCA intermediates were quantified using external standard curves.

Dried polar and nonpolar metabolites were processed for gas chromatography (GC) mass spectrometry (MS) as described previously in Cordes and Metallo (Cordes and Metallo, 2019). Briefly, polar metabolites were derivatized using a Gerstel MultiPurpose Sampler (MPS 2XL). Methoxime-tBDMS derivatives were formed by addition of 15 μL 2% (w/v) methoxylamine hydrochloride (MP Biomedicals, Solon, OH) in pyridine and incubated at 45°C for 60 minutes. Samples were then silylated by addition of 15 μL of N-tert-butyldimethylsily-N-methyltrifluoroacetamide (MTBSTFA) with 1% tert-butyldimethylchlorosilane (tBDMS) (Regis Technologies, Morton Grove, IL) and incubated at 45°C for 30 minutes. Nonpolar metabolites were saponified and transesterified to fatty acid methyl esters (FAMEs) by adding 500 µL of 2% H_2_SO_4_ in methanol to the dried nonpolar layer and heating at 50°C for 1 hour. FAMEs were then extracted by adding 100 µL of a saturated salt solution and 500 µL hexane and vortexing for 1 minute. The hexane layer was removed, evaporated, and resuspended with 60µL hexane for injection.

Derivatized polar samples were injected into a GC-MS using a DB-35MS column (30m x 0.25mm i.d. x 0.25μm, Agilent J&W Scientific, Santa Clara, CA) installed in an Agilent 7890B GC system integrated with an Agilent 5977a MS. Samples were injected at a GC oven temperature of 100°C which was held for 1 minute before ramping to 255°C at 3.5°C/min then to 320°C at 15°C/min and held for 3 minutes. Electron impact ionization was performed with the MS scanning over the range of 100-650 m/z for polar metabolites.

Derivatized nonpolar samples were injected into a GC-MS using a Fame Select column (100m x 0.25mm i.d. x 0.25μm, Agilent J&W Scientific, Santa Clara, CA) installed in an Agilent 7890A GC system integrated with an Agilent 5977A MS. Samples were injected at a GC oven temperature of 80°C which was held for 1 minute before ramping to 170°C at 20°C/min then to 188°C at 1°C/min then to 250°C at 20°C/min and held for 10 minutes. Electron impact ionization was performed with the MS scanning over the range of 54-400 m/z for nonpolar metabolites.

Metabolite levels and mass isotopomer distributions were analyzed with an in house Matlab script which integrated the metabolite fragment ions and corrected for natural isotope abundances. Mole percent enrichment calculations were performed using Escher-Trace (Kumar et al., 2020).

#### RNA isolation and quantitative RT-PCR

Total RNA was purified from cultured cells using Trizol Reagent (Life Technologies) per manufacturer’s instructions. First-strand cDNA was synthesized from 1 μg of total RNA using iScript Reverse Transcription Supermix for RT-PCR (Bio-Rad Laboratories) according to the manufacturer’s instructions. Individual 20 μl SYBR Green real-time PCR reactions consisted of 1 μl of diluted cDNA, 10 μl of SYBR Green Supermix (Bio-Rad), and 1 μl of each 5 μM forward and reverse primers. For standardization of quantification, 18S was amplified simultaneously. The PCR was carried out on 96-well plates on a CFX Connect Real time System (Bio-Rad), using a three-stage program provided by the manufacturer: 95 °C for 3 min, 40 cycles of 95 °C for 10 s and 60 °C for 30 s. Gene-specific primers are tabulated in Table S1.

#### ^14^C citrate uptake assay

Uptake of ^14^C citrate was quantified as described previously (Pajor and Sun, 2010). Briefly, cells were seeded at 3 x 10^5^ cells/well on 6 well collagen treated plates. After 24 hours, cells were acclimated to serum free media for 24 hours. To carry out the assay, each well was washed twice with choline buffer (mM: 140 Choline, 2 KCl, 1 MgCl_2_, 1 CaCl_2_, 10 HEPES, pH adjusted to 7.4 with 1M Tris), then incubated with 1 ml sodium buffer (mM: 140 NaCl, 2 KCl, 1 MgCl_2_, 1 CaCl_2_, 10 HEPES, pH adjusted to 7.4 with 1M Tris) or choline buffer containing 100 µM [^14^C]-citrate for 30 minutes at 37°C with rocking. The uptake assays were stopped, and the cells were washed four times with 2.5 ml choline buffer. Cells were dissolved in 0.4 ml/well 1% SDS, transferred to scintillation vials with Econoscint Scintillation cocktail and then the radioactivity in the plates was counted using a scintillation counter.

### Quantification and Statistical Analysis

Statistical analyses were performed using Graphpad PRISM. Unless indicated, all results shown as mean ± SD of cellular triplicates obtained from one representative experiment as specified in each figure legend. P values were calculated using a Student’s two-tailed *t* test, One-way ANOVA w/ Dunnet’s method for multiple comparisons, or Two-way ANOVA w/ Tukey’s method for multiple comparisons; *, P value < 0.05; **, P value < 0.01; ***, P value < 0.001, ****, P value < 0.0001. Unless indicated, all normalization and statistical tests compared to NTC cells, normoxia, or (-)citrate conditions.

## Supplemental Information

Document S1: Figures S1-S4, Table S1

