## Supplementary material for "NaCT (*SLC13A5*) facilitates citrate import and metabolism under nutrient-limited conditions": Document S1: Figures S1-S4, Table S1

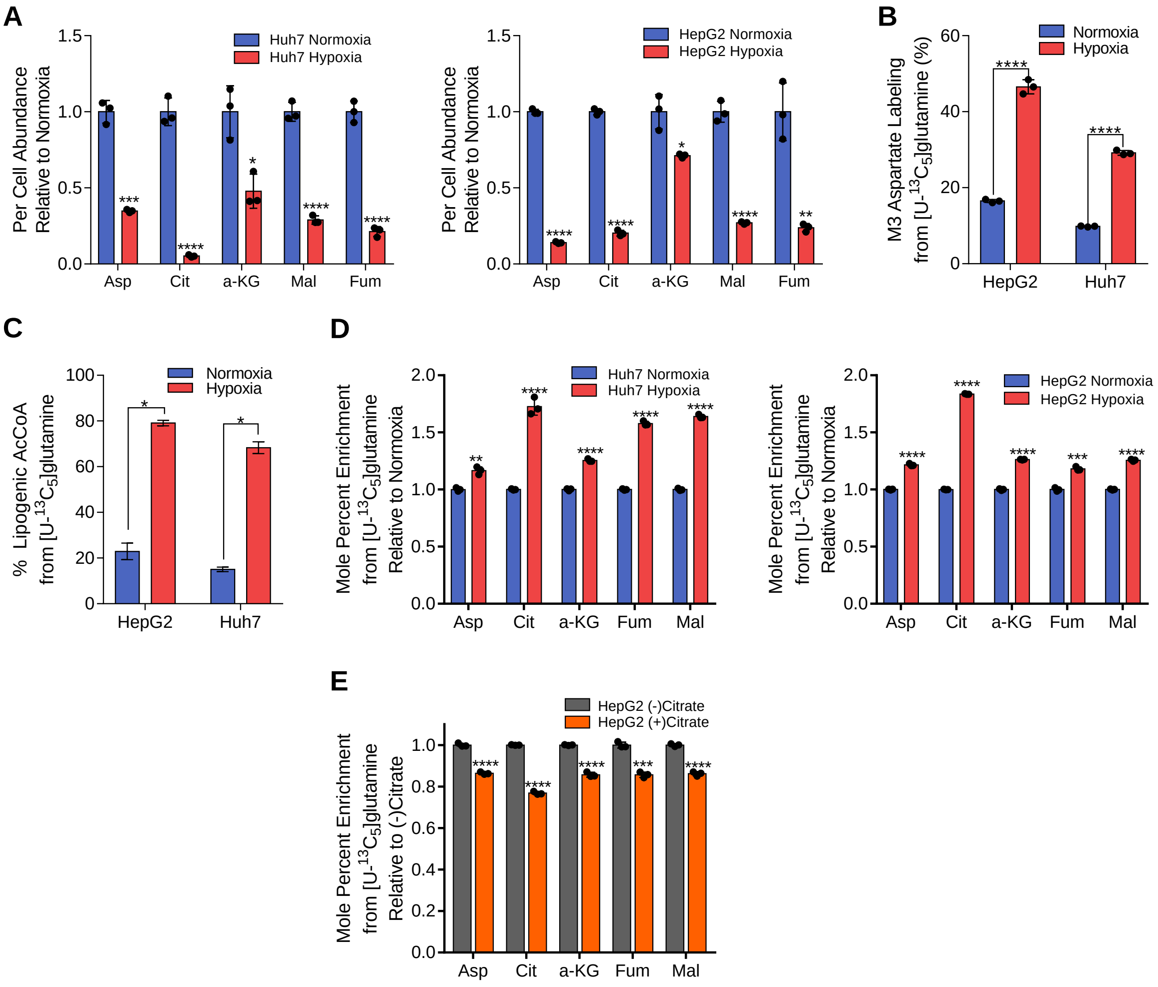


**Supplementary Figure 1. Citrate dilutes central carbon pathway labeling in hepatocellular carcinoma and neuronal cells in hypoxia. Related to Figure 2.**

(A) Per cell abundances of TCA intermediates in Huh7 (left) and HepG2 (right) cells grown in normoxia or hypoxia for 48 hours, relative to normoxia (n=3).

(B) Percent labeling of M3 aspartate from [U-$^{13}C_5$]glutamine in HepG2 and Huh7 cells grown in normoxia or hypoxia for 48 hours (n=3).

(C) Percent of lipogenic acetyl-CoA contributed by [U-$^{13}C_5$]glutamine in HepG2 and Huh7 cells grown in normoxia or hypoxia for 48 hours (n=3).

(D) Mole percent enrichment of TCA intermediates from [U-$^{13}C_5$]glutamine in Huh7 (left) and HepG2 (right) cells grown in normoxia or hypoxia for 48 hours, relative to normoxia (n=3).

(E) Mole percent enrichment of TCA intermediates from [U-$^{13}C_5$]glutamine in HepG2 cells grown in hypoxia +/- 500 μM citrate for 48 hours, relative to (-) citrate (n=3).

In (A,B,D,E) data are plotted as mean ± SD. Statistical significance is relative to normoxia (A,B,D) or (-) citrate (E) as determined by two-sided Student's t-test with *, P value < 0.05; **, P value < 0.01; ***, P value < 0.001, ****, P value < 0.0001. In (C) data are plotted as mean ± 95% confidence interval (CI). Statistical significance by non-overlapping confidence intervals, *. Unless indicated, all data represent biological triplicates. Data shown are from one of at least two separate experiments.

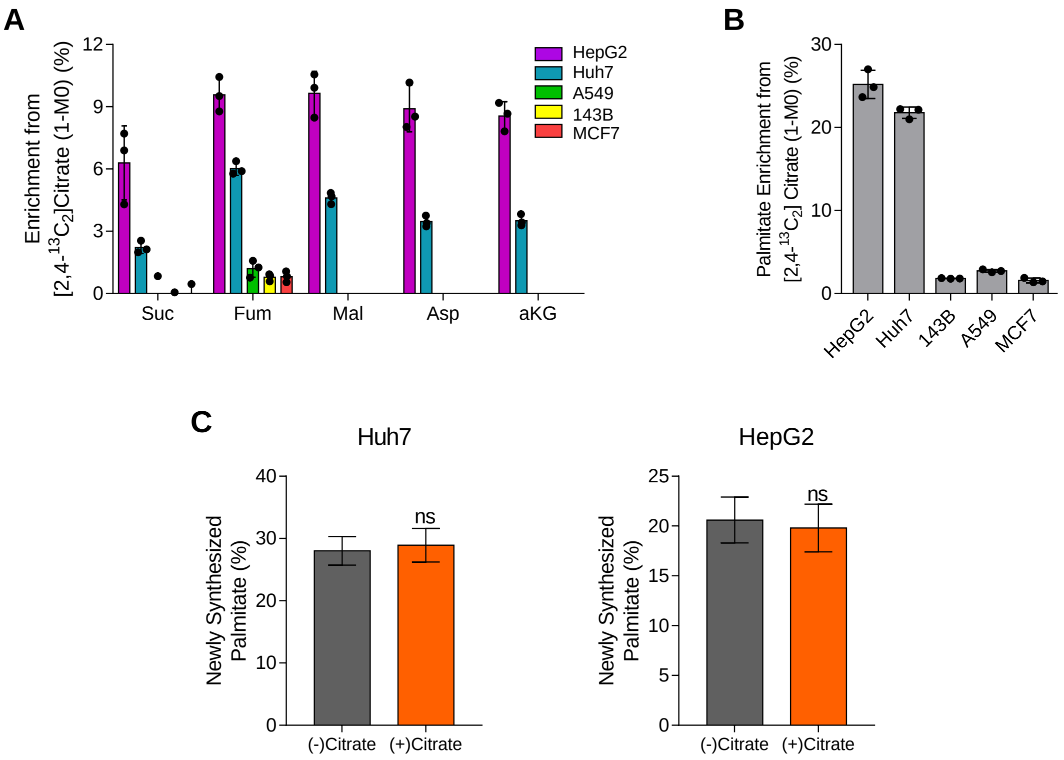


**Supplemental Figure 2. Exogenous citrate is metabolized for TCA anaplerosis and fatty acid synthesis. Related to Figure 3.**

(A) Enrichment (1-M0) of TCA intermediates from 500 µM [2,4-$^{13}C_2$]citrate in cancer cells over 48 hours in hypoxia (n=3).

(B) Palmitate enrichment (1-M0) from 500 µM [2,4-$^{13}C_2$]citrate in cancer cells over 48 hours in hypoxia (n=3).

(C) De novo synthesis of palmitate +/- 500 µM citrate in Huh7 (left) and HepG2 (right) cells grown in hypoxia over 48 hours

In (A,B) data are plotted as mean ± SD. Unless indicated, all data represent biological triplicates. In (C) data are plotted as mean ± 95% confidence interval (CI). Statistical significance by non-overlapping confidence intervals, *. Data shown are from one of at least two separate experiments.

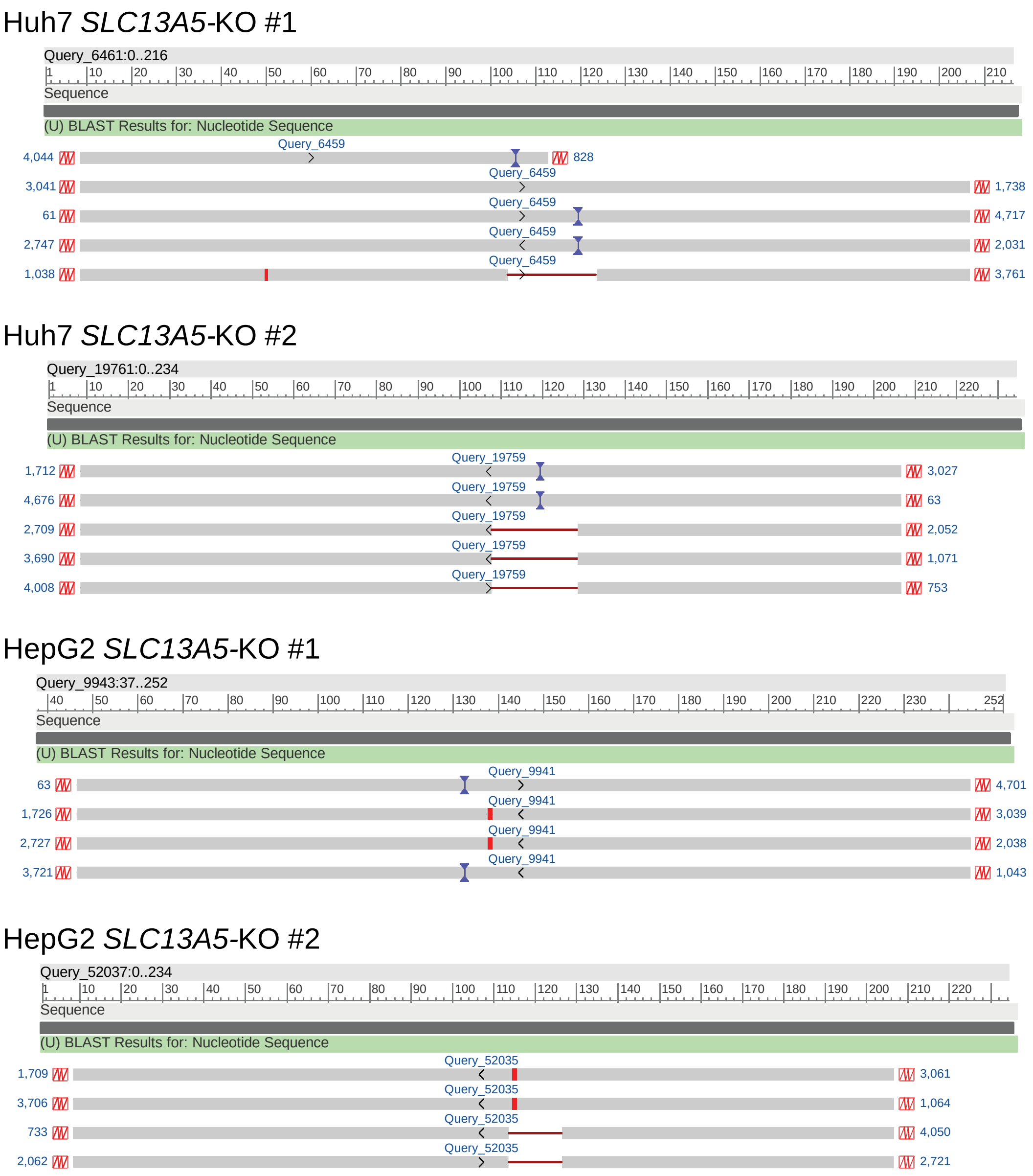


**Supplemental Figure 3. Alignment of DNA sequences obtained from CRISPRcas9 HCC *SLC13A5*-KO clones. Related to Figure 5.**

Sequences aligned using NCBI BLASTN suite (Agarwala et al., 2018). Results were visualized using NCBI Viewer 3.41.1.

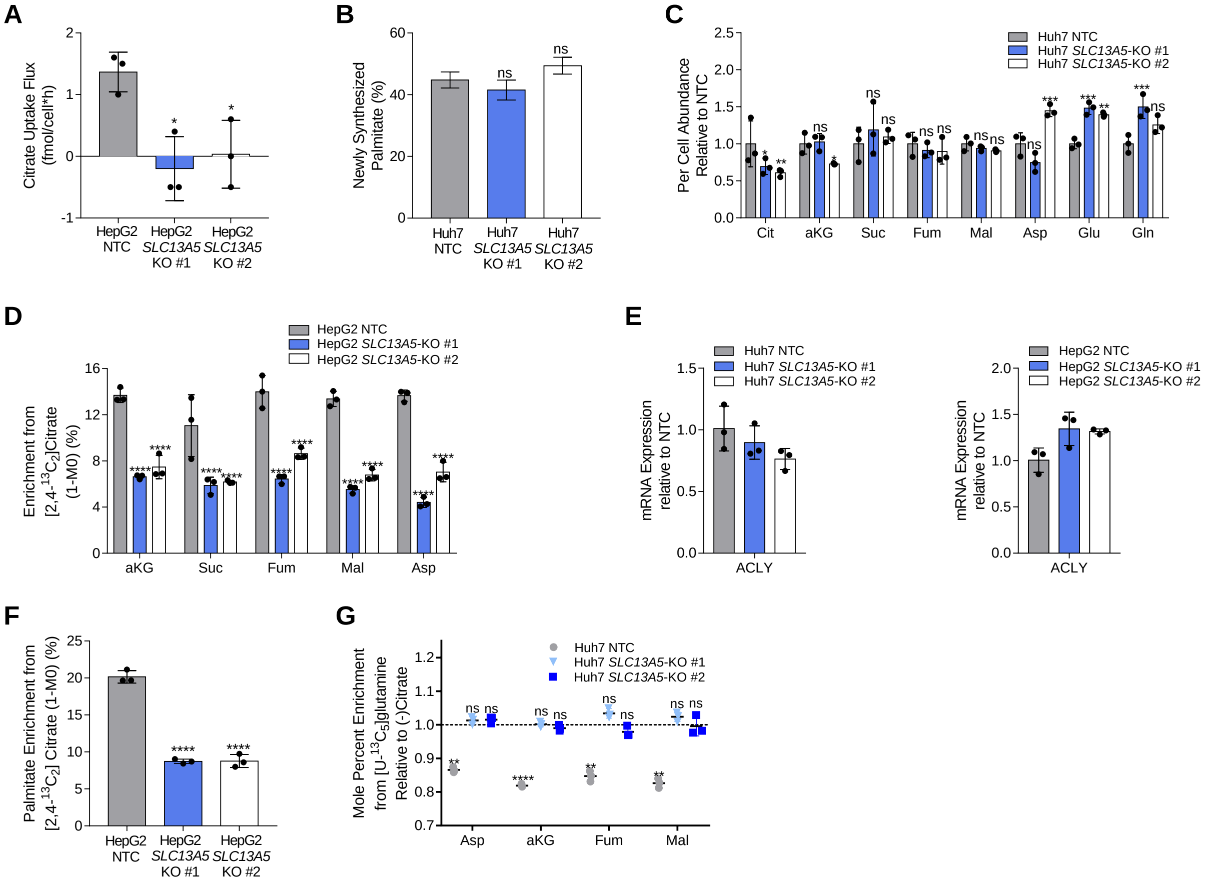


**Supplemental Figure 4. NaCT coordinates citrate import and metabolism in hepatocellular carcinoma cells. Related to Figure 5.**

(A) Citrate uptake flux over 48 hours in hypoxia in HepG2 NTC and *SLC13A5*-KO cells (n=3).

(B) De novo synthesis of palmitate in Huh7 NTC and *SLC13A5*-KO cells grown in hypoxia for 48 hours (n=3).

(C) Per cell abundance of metabolites in Huh7 NTC and *SLC13A5*-KO cells grown in hypoxia for 48 hours, relative to NTC (n=3).

(D) ACLY mRNA expression in NTC and *SLC13A5*-KO Huh7 (left) and HepG2 (right) cells grown in normoxia, relative to NTC (n=3).

(E) Enrichment (1-M0) of TCA intermediates from [2,4-$^{13}C_2$]citrate in HepG2 NTC and *SLC13A5*-KO cells grown in hypoxia for 48 hours (n=3).

(F) Enrichment (1-M0) of palmitate from [2,4-$^{13}C_2$]citrate in HepG2 NTC and *SLC13A5*-KO cells grown in hypoxia for 48 hours (n=3).

(G) Mole percent enrichment of TCA intermediates from [U-$^{13}C_5$]glutamine in Huh7 NTC and *SLC13A5*-KO cells +/- 500 µM citrate grown in hypoxia for 48 hours, relative to (-) citrate (n=3).

In (A,C-F) all graphs data are plotted as mean ± SD. Statistical significance is relative to NTC as determined by One-way ANOVA w/ Dunnet's method for multiple comparisons (A,C-F) or relative to (-) citrate as determined by two-sided Student's t-test (G) with *, P value < 0.05; **, P value < 0.01; ***, P value < 0.001, ****, P value < 0.0001. In (B) data are plotted as mean ± 95% confidence interval (CI). Statistical significance by non-overlapping confidence intervals, *. Unless indicated, all data represent biological triplicates. Data shown are from one of at least two separate experiments.

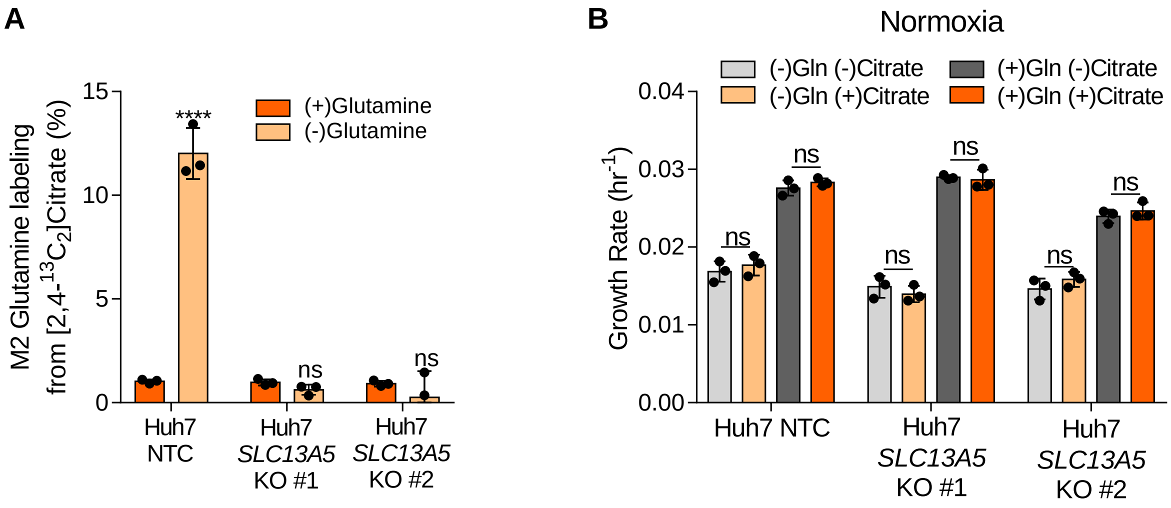


**Supplemental Figure 5. NaCT facilitates growth under nutrient stress and resistance to zinc toxicity. Related to Figure 6.**

(A) Mole percent enrichment of glutamine from 500 μM [2,4-$^{13}C_2$]citrate in Huh7 NTC and *SLC13A5*-KO cells over 48 hours in hypoxia (n=3).

(B) Growth rates of Huh7 NTC and *SLC13A5*-KO cells grown in high glucose DMEM +/- 4mM glutamine +/- 500 μM citrate in normoxia for 4 days (n=3).

In all graphs data are plotted as mean ± SD. Statistical significance is determined by two-sided Student's t-test relative to (+) glutamine (A); or determined by Two-way ANOVA w/ Tukey's method for multiple comparisons relative to (-) citrate (B) with *, P value < 0.05; **, P value < 0.01; ***, P value < 0.001, ****, P value < 0.0001. Unless indicated, all data represent biological triplicates. Data shown are from one of at least two separate experiments.

**Table S1. Oligonucleotide sequences used in this study**

| Primer Name | Sequence | Application |
|---|---|---|
| ACLY (human) fwd | TCGGCCAAGGCAATTTCAGAG | qRT-PCR |
| ACLY (human) rev | CGAGCATACTTGAACCGATTCT | qRT-PCR |
| SLC13A5 (human) fwd | CTGCCACTCGTCATTCTGATG | qRT-PCR |
| SLC13A5 (human) rev | ATGTTGGTGTCCTTCATGTACTG | qRT-PCR |
| r18s (human) fwd | AGTCCCTGCCCTTTGTACACA | qRT-PCR |
| r18s (human) rev | CGATCCGAGGGCCTCACTA | qRT-PCR |
| Slc13a5 (rat) fwd | GGTGCACAGATGTCATCCCA | qRT-PCR |
| Slc13a5 (rat) rev | AGCATGTTGGTGTCCGTCAT | qRT-PCR |
| r18s (rat) fwd | AGAAACGGCTACCACATCCA | qRT-PCR |
| r18s (rat) rev | CTCGAAAGAGTCCTGTATTGT | qRT-PCR |
| SLC13A5 (human) PCR fwd | AGGCATCCCATAGTGACCCT | Target Site PCR Primer |
| SLC13A5 (human) PCR rev | CACAGAACTGCCGGAGTTGT | Target Site PCR Primer |
| sgRNA-NTC-fwd | GGCCGTGTTGCTGGATACGCC | CRISPR/Cas9 |
| sgRNA-NTC-rev | GGCGTATCCAGCAACACGGCC | CRISPR/Cas9 |
| sgRNA-SLC13A5-fwd | AGGCACAATGAATAGCAGGG | CRISPR/Cas9 |
| sgRNA-SLC13A5-rev | CCCTGCTATTCATTGTGCCT | CRISPR/Cas9 |
